# A mathematical model of ephaptic interactions in neuronal fiber pathways: could there be more than transmission along the tracts?

**DOI:** 10.1101/698282

**Authors:** Hiba Sheheitli, Viktor K. Jirsa

## Abstract

In the past several decades, there has been numerous experimental and modeling efforts to study ephaptic interactions in neuronal systems. While studies on the matter have looked at either axons of the peripheral nervous system or cortical neuronal structures, no attention has be given to the possibility of ephaptic interactions in the white matter tracts of the brain. Inspired by the highly organized and tightly packed geometry of axons in neuronal fiber pathways, we aim to theoretically investigate the potential effects of ephaptic interactions along these structures that are resilient to experimental probing. For that end, we use axonal cable theory to derive a minimal model of a sheet of N ephaptically coupled axons. We numerically solve the equations and explore the dynamics of the system as the ephaptic coupling parameter is varied. We demonstrate that ephaptic interactions can lead to local phase locking between impulses traveling along adjacent axons. As ephaptic coupling is increased, traveling impulses trigger new impulses along adjacent axons resulting in finite size traveling fronts. For strong enough coupling, impulses propagate laterally and backwards, resulting in complex spatio-temporal patterns. While it is common for large scale brain network models to assume the role of brain fiber pathways to be that of mere transmission of signals between different brain regions, our work calls for a closer re-examination of the validity of such a view. The results suggest that in the presence of significant ephaptic interactions the brain fiber tracts can act as a dynamic active medium.

**Author summary:** Starting from local circuit theory and the Fitzhugh-Nagumo cable model of an axon, we derive a system of nonlinear coupled partial differential equations (PDE’s) to model a sheet of N ephaptically coupled axons. We also put forward a continuous limit approximation that transforms the model into a field equation in the form of a two-dimensional PDE that allows for the extension of the model to a 3D domain. We numerically solve the equations and explore the dynamic responses as the ephaptic coupling strength is varied. We observe that ephaptic interaction allows for phase locking of adjacent impulses and coordination of subthreshold dynamics. In addition, when strong enough, ephaptic interaction can lead to the generation of new impulses along the axons as well as lateral and backward propagation in the form of traveling fronts and complex spatio-temporal patterns. The transition between different dynamic regimes happens abruptly at critical values of the parameter. We also compare the dynamics of the two models and find good qualitative correspondence in certain parameter regimes. The results put into question the validity of assuming the role of fiber pathways to be that of mere interneuronal transmission and calls for further investigation of the matter.

## Introduction

It has been long thought that signals exchanged between different brain regions are faithfully transmitted along the white matter tracts through axons that can be modelled as passive electric cables [1]. This has led many large scale network models to assume that signals communicated between differetnt brain regions are relayed along the axons of fiber pathways with a finite speed without any interaction occurring between the traveling signals along the way [2–4]. The aim of this work is to motivate a re-examination of this latter highly consequential assumption. In 1940, Katz & Schmitt [5] investigated the non-synaptic electrical interaction between adjacent nerve fibers. In their work, two large parallel non-myelinated axons were isolated from the crab limb nerve. They succeeded in demonstrating that: 1) the passage of an action potential (impulse) in one fiber causes subthreshold excitability changes in the adjacent fiber and 2) when impulses are set up simultaneously along both fibers, a mutual interaction occurs that can lead to speeding up or slowing down of the impulses and also possibly to synchronization between the two impulses, depending on the initial phase relationship. The effect was observed to be amplified when the resistance of the extracellular space surrounding the axons was increased. In the next year, a similar study was presented by Arvanitaki [6] on giant axons of Sepia officinalis (common cuttlefish). In that study, Arvanitaki coined the term “ephapse” to denote “the locus of contact or close vicinity of two active functional surfaces”. The term is derived from the Greek term signifying the act of touching, as opposed to “synapse” which is derived from the Greek term signifying the act of joining or linking. Since then, the term ephaptic interaction has been used to refer to communication between neuronal cells via electrical conduction through the surrounding extracellular space, as opposed to communication mediated by chemical synapses or gap-junctions. In 1980, ephaptic transmission was observed between spontaneously active single nerve fibers in the spinal nerve roots of dystrophic mice [7]. Shortly after, ephaptic interactions were observed to contribute to neuronal synchrony in rat hippocampal slices [8]. Then in 1984, experiments suggested a role for ephaptic transmission in hemifacial spasm pathophysiology by causing “cross-talk” between facial nerve fibers [9]. More recently, hallmarks of ephaptic interaction were observed in rat cortical pyramidal neurons in slices, and supported the idea that this interaction facilitates the coordination and possibly the synchrony of neighboring neurons in the gray matter [10]. In addition, there has been numerous other experimental and modeling investigations of ephaptic interaction [11–19]. However, it can be seen that all these previous studies focused on one of two contexts: 1) cortical areas, particularly interactions between neighboring neurons through the resulting local field potential [8, 10, 15, 17, 19–21] or 2) peripheral nerves, particularly interactions between myelinated axons in a nerve bundle and inquiries into effects of demyelination [7, 9, 11, 22–25]. To our knowledge, there has been no discussion on ephaptic interaction between axons of the white matter tracts. While the predominant myelination in white matter axons might be presumed to be preventing ephaptic interference, studies of myelinated axons in nerves suggest otherwise [22, 24, 26]. Moreover, in some fiber pathways the proportion of unmyelinated axons can be as high as 30% [27]. It is known that fiber pathways in the brain are constituted of densely packed long axons running in parallel in what can be seen as two dimensional sheets [28]. Electron micrograph images show that neighboring axons in fiber pathways are often separated by distances as small as 0.1nm [29] which would suggest a relatively high extracellular space resistance. These latter geometric characteristics set favorable conditions for ephaptic exchanges to be at play in white matter fiber pathways. Ideally, direct experimental examination of the activity of axons in the white matter would serve to accurately quantify ephaptic interactions there. However, probing into the inner workings of the white matter remains a challenging endeavor, mainly due to technical limitations on the temporal and spatial resolution of current noninvasive imaging techniques [30]. Inspired by these facts, we wish to investigate the matter by putting forward a simple but realistic mathematical model of excitable axons arranged in a sheet like geometry and coupled through a resistive extracellular space.

In section 1, we start from local circuit theory and the cable model of an axon to derive a model for a sheet of N ephaptically coupled axons. We then make a continuous limit approximation to transform the resulting model of N coupled 1D PDE’s into a 2D PDE that can be seen as a field equation governing the dynamics of a sheet of coupled axons. In section 2, we numerically solve the equations and explore the different possible dynamical regimes along with examining the equivalence of the two proposed models. In section 3, we discuss the potential ramifications of the results along with future work directions that this work motivates.

## 1 Materials and methods

### 1.1 The mathematical model

Our goal here is to put forward a minimal model that possesses the key elements that allow the study of the effects of ephaptic interactions on action potential transmission along fiber pathways. Given the densely packed parallel geometric arrangement of axons in the white matter, we assume that currents generated during action potential propagation are mainly axial in direction, both inside the axons and in the surrounding extracellular space [31, 32]. Then the axons can be represented by what’s known as the cable equation while the extracellular space between axons can be represented by an effective longitudinal resistance per unit length [33, 34]. Such a model for two ephaptically coupled axons is derived in [12]. Fig.1 depicts the equivalent circuit model used to derive the cable equations for two ephaptically coupled axons. The following notation is used here:

*i*^*a*^: axial (axoplasmic) current inside the axon per unit length

*i*^*e*^: axial current in the extracellular space surrounding the axons per unit length

*i*^*m*^: axonal transmembrane current per unit area

*r*_*a*_: axoplasmic resistance per unit length

*r*_*e*_: extracellular space resistance per unit length

*c*: axonal membrane capacitance per unit length

*j*_*ion*_: active ionic current flowing across the axonal membrane per unit length

*g*: membrane conductance per unit length

*z*: distance along the axon

*v*^*m*^: transmembrane potential of an axon

*v*^*a*^: axoplasmic potential inside an axon

*v*^*e*^: extracellular space potential

*I*: external applied current per unit length

*a, b*: parameters of the Fitzhugh-Nagumo model

**Fig 1.**
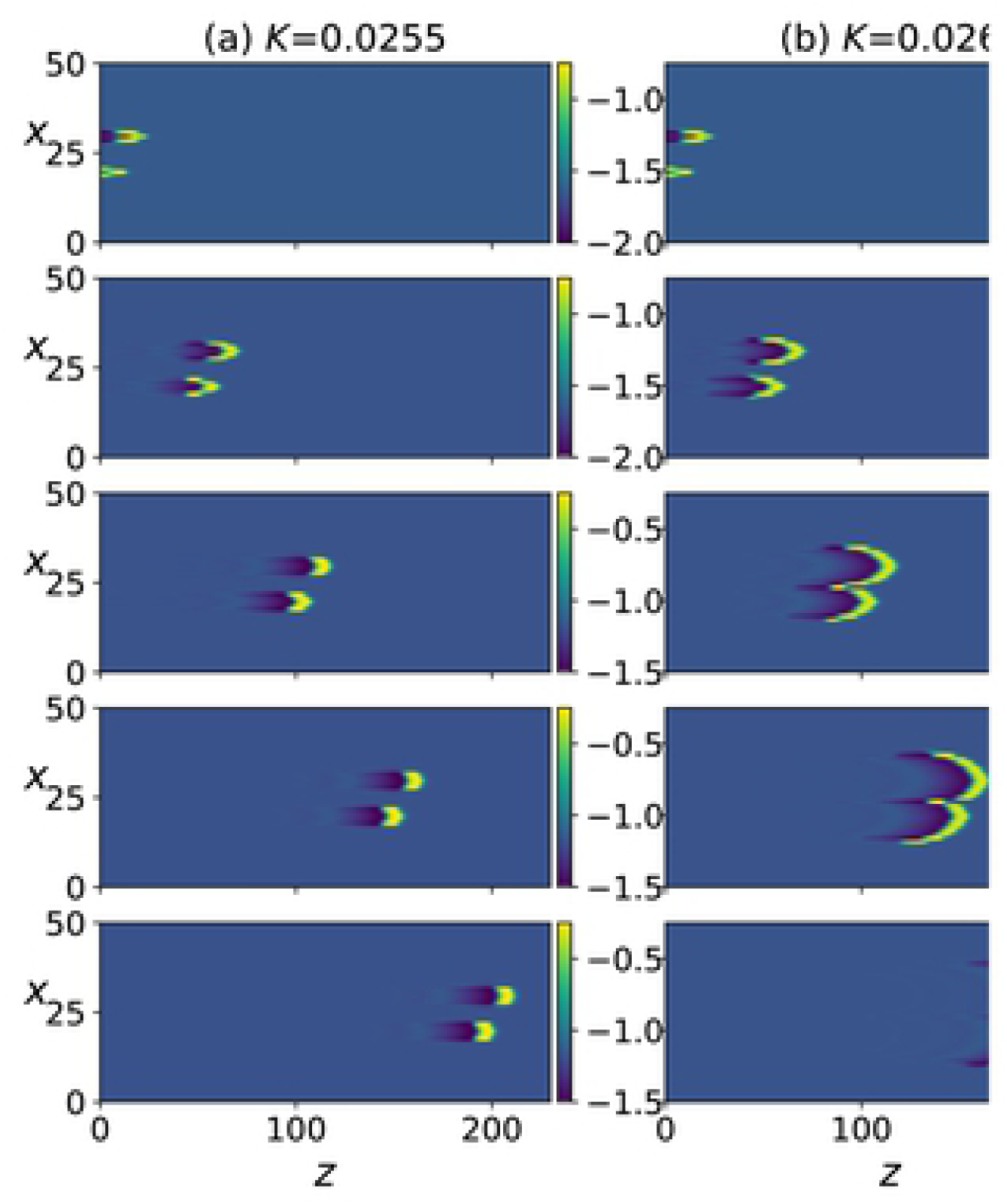
Schematic of the equivalent circuit model for two ephaptically coupled axons

Here, all currents and potential variables are varying functions of time and axial location *z*.

In the limit of Δ*z* → 0, Kirchoff’s current law gives the following relationships between the transmembrane, axial and extracellular currents; subscripts 1 and 2 each refer to one of the two identical axons:

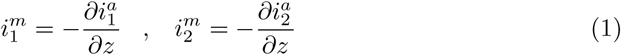

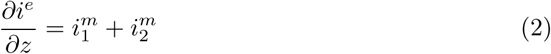

In addition, Ohm’s law relates the currents to the electric potentials as follows:

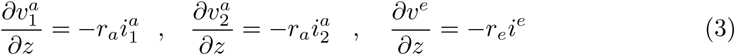

Furthermore, the transmembrane current for each axon can be expressed as:

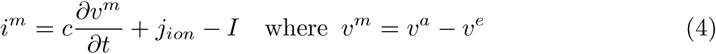

The term *j*_*ion*_ represents the active transmembrane currents due to ion channels activity which is nonlinearly dependent on the transmembrane voltage. Detailed mathematical representation of the dependence of *j*_*ion*_ on *v*^*m*^ was described in the seminal work by Hodgkin and Huxley [1], which utilized three variables to represent the kinetics of ion channels activation. In 1961, Fitzhugh proposed a simplification of that model which utilizes only one recovery variable [35]:

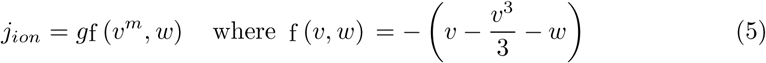

Here, *w* is a slow recovery variable. To arrive at the cable equation model, we need to combine all the above relationships to eliminate the current variables. We start by differentiating the expression for *v*^*m*^ with respect to *z* and substituting Eqs.3 in it:

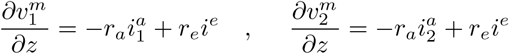

Differentiating again and substituting Eqs.1, we obtain:

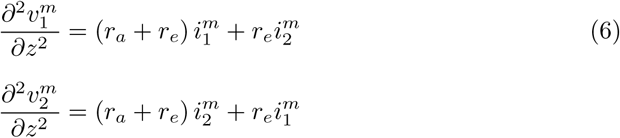

Solving the above system of two equations for an expression for 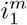 and 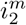, then combining the result with Eq.4, we arrive at the cable equations for two ephaptically coupled axons:

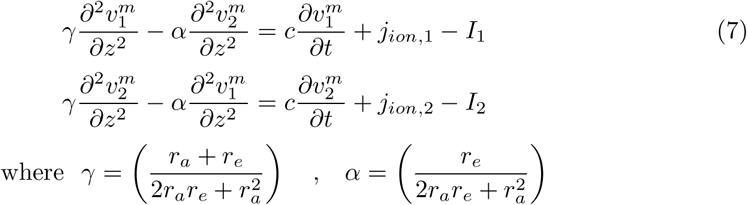

We can see that for zero extracellular resistance, the two cable equations are uncoupled such that any current exiting one axon will immediately dissipate in the extracellular space and no exchange between the axons can occur. The resulting single Fitzhugh-Nagumo cable was first put forward by Nagumo in [36]. It can be seen that the cable equation is the classical 1D diffusion equation with an added term, *j*_*ion*_. The presence of the nonlinear active currents renders the cable excitable, such that, if the membrane potential is perturbed from is resting value, it will return to that value unless the perturbation is strong enough to elicit the large action potential response which will then be propagated along the axon, away from the location of perturbation, with the signal’s shape preserved.

We wish to extend the model to a sheet of ephaptically coupled axons, that is, an *N* number of axons coupled through the extracellular space. Fig.2 shows a schematic of a cross-sectional view of such an arrangement where we represent the cross-sections of axonal cables as nodes on a line, inter-spaced with nodes representing extracellular space.

**Fig 2.**
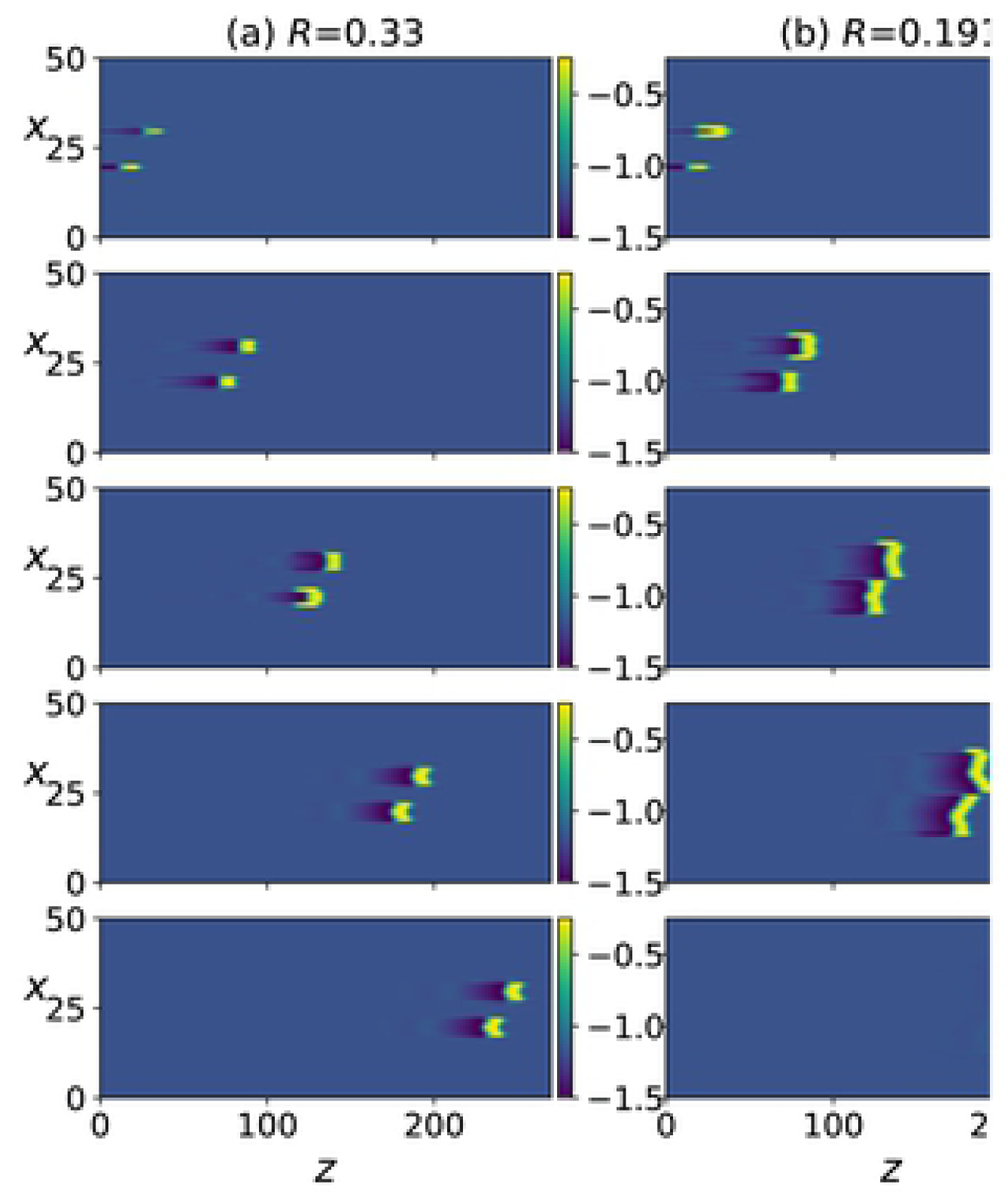
Schematic of a cross-sectional view of the sheet of N axons model.

A model of such a configuration of N number of coupled axons was presented in [13]. The main assumption made in this latter work is that each axon is only coupled to the two axons that are positioned directly next to it. While this later assumption is common in networks models, the authors offered no physiological justification for it in the context of ephaptically coupled axons. Instead, we will start from the more physical assumption that transmembrane currents are radially uniform, such that we can reasonably consider *i*^*m*^ for each cable to be equally partitioned into two parts feeding into the extracellular space (represented as nodes) adjacent to it. Consequently, while the previously presented model restricts ephaptic interactions to nearest neighbors axons [13], our model allows each axon to interact with all other axons through the shared extracellular space with the coupling strength decaying with distance between interacting axons. To arrive to that, we take the extracellular space potential for an axon positioned at a node *q* to be the average of the potential at its adjacent extracellular nodes such that:

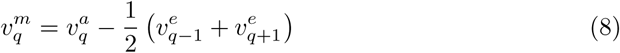

Note that the index *q* refers to the node number on the line, so for *N* axons, *q* takes values between 1 and 2*N* + 1, such that 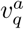 is defined on nodes *q* = 2, 4, *…, N* and 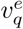 on nodes *q* = 1, 3, *…, N* + 1. Differentiating Eq.8 with respect to *z*, and making use of Ohm’s law, we obtain:

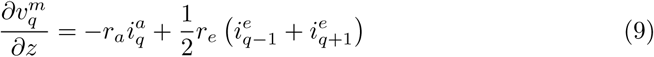

From Kirchoff’s first law of current, we have the following:

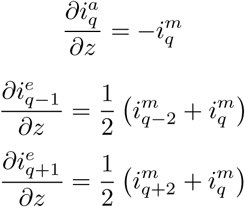

Differentiating Eq.9 again and plugging in the above current relationships, we arrive at:

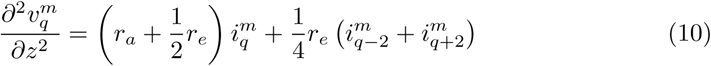

For the cables that are at the two ends of the line, the corresponding relationship would be:

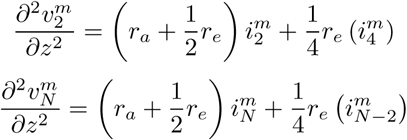

We’ve obtained a system of N equations relating *i*^*m*^ and *v*^*m*^ of all the axons. This is the equivalent of Eqs.6 for the two axon system. The linear system of N equations can be solved, such that we can express each *i*^*m*^ explicitly in terms of *v*^*m*^ of all the axons. The solution takes the form:

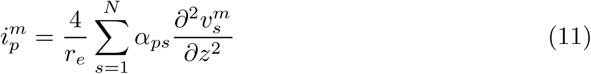

*p* = 1, 2, …, *N* referring to the N axons. The *α*’s represent coupling strength between each pair of axons and are obtained as the elements of the inverse matrix *A*^-1^ = [*α*_*ij*_] where *A* is the tridiagonal matrix:

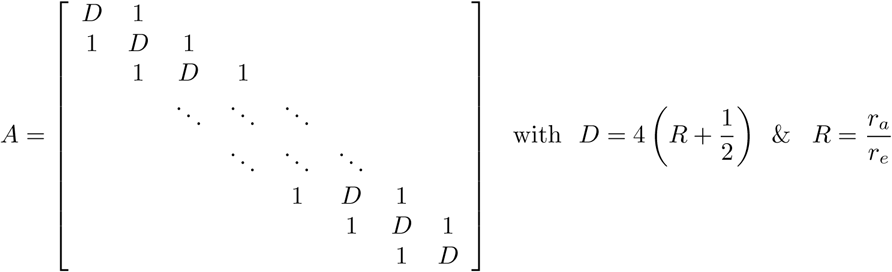

Explicit algebraic expressions for the elements of the inverse of such a tridiagonal matrix are presented in [37] and indicate that the ephaptic effect increases as the ratio *R* decreases and that *α* for two axons on the line decreases as the distance between them increases. However, for the numerical solutions presented in the sections to follow, we found it simpler to numerically compute the inverse of *A* instead of the individual *α*′ *s*.

Combining Eqs.11, 5 and 4, we obtain the model for a sheet of N ephaptically coupled Fitzhugh-Nagumo cables:

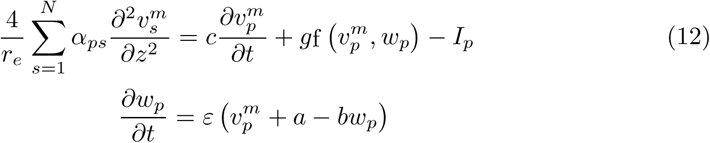

We non-dimensionalize the space and time variables as follows:

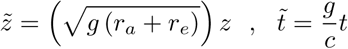

The system becomes:

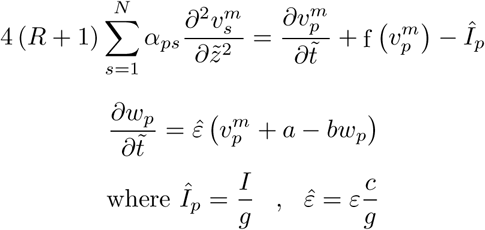

From now on, we drop the superscript of the transmembrane voltage along with the tilde and hat. The final equations take the form:

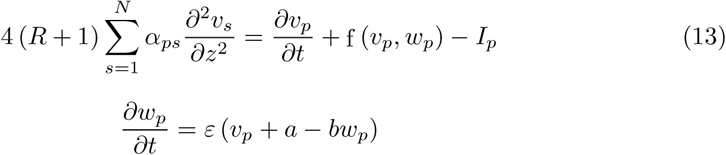

Without loss of generality, for the results that will follow, we choose the following values for the Fitzhugh-Nagumo recovery variable: *a* = 0.7, *b* = 0.5 and *E* = 0.1. Then, we are left with one free parameter *R* which reflects the strength of the ephaptic interaction. The goal is to investigate the dynamics of the system as this parameter is varied.

### 1.2 Estimation of the coupling strength parameter

To estimate the physically plausible range of values for parameter *R*, we start from the definition:

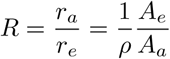

where *A*_*e*_ and *A*_*a*_ are the cross-sectional areas of the extracellular space and the axon, respectively. *ρ* is the ratio of extracellular to intracellular resistivity, and is typically assumed to be in the range of 1 to 4 [19]. Given that axons of the fiber pathways are very tightly packed, we consider that the cross-sectional area of the space between adjacent axons can range from a tenth to several multiples of the cross-sectional area of the axon. Based on that, we consider *R* to vary between 0 and 1.

### 1.3 Continuous limit approximation

Our model is a system of N coupled nonlinear PDE’s each representing one distinct cable and accompanied by an ordinary differential equation for the corresponding slow recovery variable. However, neighboring axons in fiber pathways are often very densely packed such as the distances separating two adjacent axons are considerably small relative to the axonal diameter [29]. For this reason, we will make the approximation that the variables *v*^*m*^ and *i*^*m*^, while being only physically defined for the axonal space, can be abstractly represented by continuous field variables *v* and *i*. If we go back to Eq.10, we notice that the last two terms can be rewritten by using the following discrete approximation of a second partial derivative [38]:

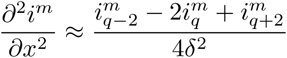

then

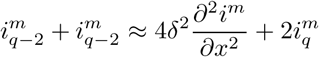

where *δ* is the small separation between adjacent axons. Then Eq.10 is transformed to:

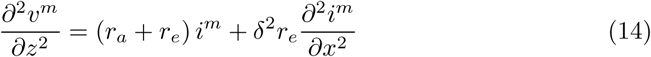

This equation relates the transmembrane voltage and current approximate field variables. The second relationship between the two is given by the balance of currents Eq.4. The approximate continuous system then takes the form:

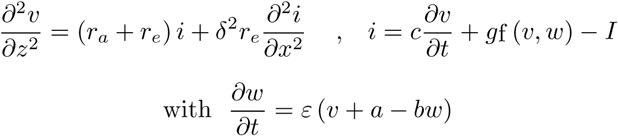

where the superscripts were dropped for brevity. Next, we nondimensionalize the equations using the following rescaling:

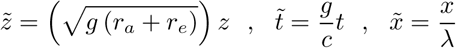

Here, *λ* is a characteristic lateral length scale of the same order of magnitude as the average axonal diameter (*µm*). The resulting system becomes:

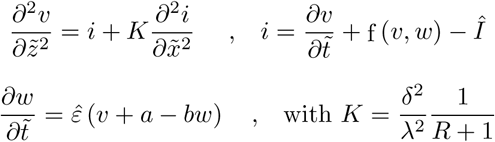

Dropping the tilde and hat for brevity, we obtain the final form of the approximate continuous field equations for a sheet of ephaptically coupled axons:

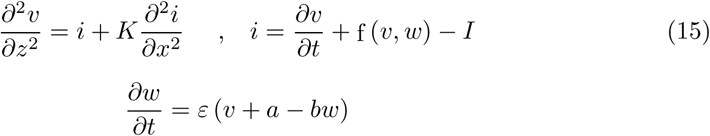

We note here that the latter equations governing *v* and *i* can be transformed into a more compact form of one partial integro–differential equation using Green’s functions and contour integration, such that the system becomes:

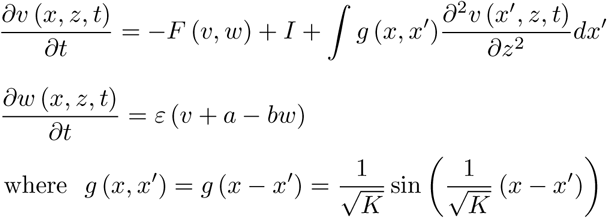

In the results that follow, we take 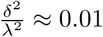, to be consistent with the assumption that the inter-axonal spacing is very small compared to the characteristic length. Hence, we consider values of *K* in the range of 0 to 0.1 to be in line with the above choice of *R* being between 0 and 1.

We also note that the model can be extended to the 3D case by considering axons on a two dimensional grid (*x, y*) instead of a 1D line (*x*). While deriving the N coupled PDE’s system will be more tedious in this case, the continuous limit approximation leads to Eqs.15 with one added term on the right hand side of the first equation:

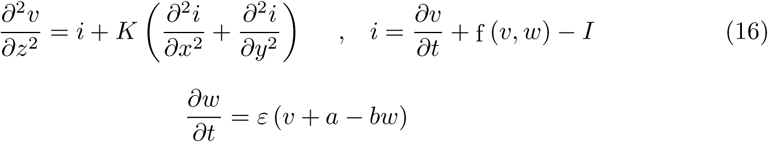

### 1.4 Numerical implementation

Numerical solutions of the two model systems were obtained using the Crank-Nicolson finite difference method. After numerical experiments were performed with smaller values and confidence in the stability of the solution was established, we chose a time step of 0.05 and a spatial step of 0.5 and 1 for the *z* and *x* directions, respectively. Zero flux boundary conditions were enforced, such that the first spatial derivative of *v* remains 0 at the boundaries for all time. To investigate the response of the system, impulses were initiated at the inlet of axons using a brief and localized input current *I* = 2 applied to the axon for *t* ∈ [0, 2] and *z* ∈ [0, 4].

## 2 Results

Numerical simulations were performed to investigate the dynamics of the system as the ephaptic interaction strength was varied. In addition, we compared the dynamics of the continuous limit system (Eqs.15) to that of the original discrete model (Eqs.13).

### 2.1 Phase interactions

In Fig.3, two adjacent axons are stimulated such that an action potential is initiated along each of them and travels from left to right. The timing of the stimulation is such that one impulse lags behind the other. The shape of the action potentials and their propagation along the *z*-direction can be seen in the supporting information (S1 Fig). In Fig.3(a), where we set *R* = 0.8 corresponding to weak ephaptic coupling, the two impulses travel independently without influencing each other. In Fig.3(b) and (c), for the same value of *R* but with the stimulated axons directly adjacent to each other, ephaptic interaction causes the two impulses to attract/repel each other such that the lag between them decreases/increases, after which they remain locked together, depending on the initial time lag between them. This type of interaction has been experimentally observed in [5].

**Fig 3.**
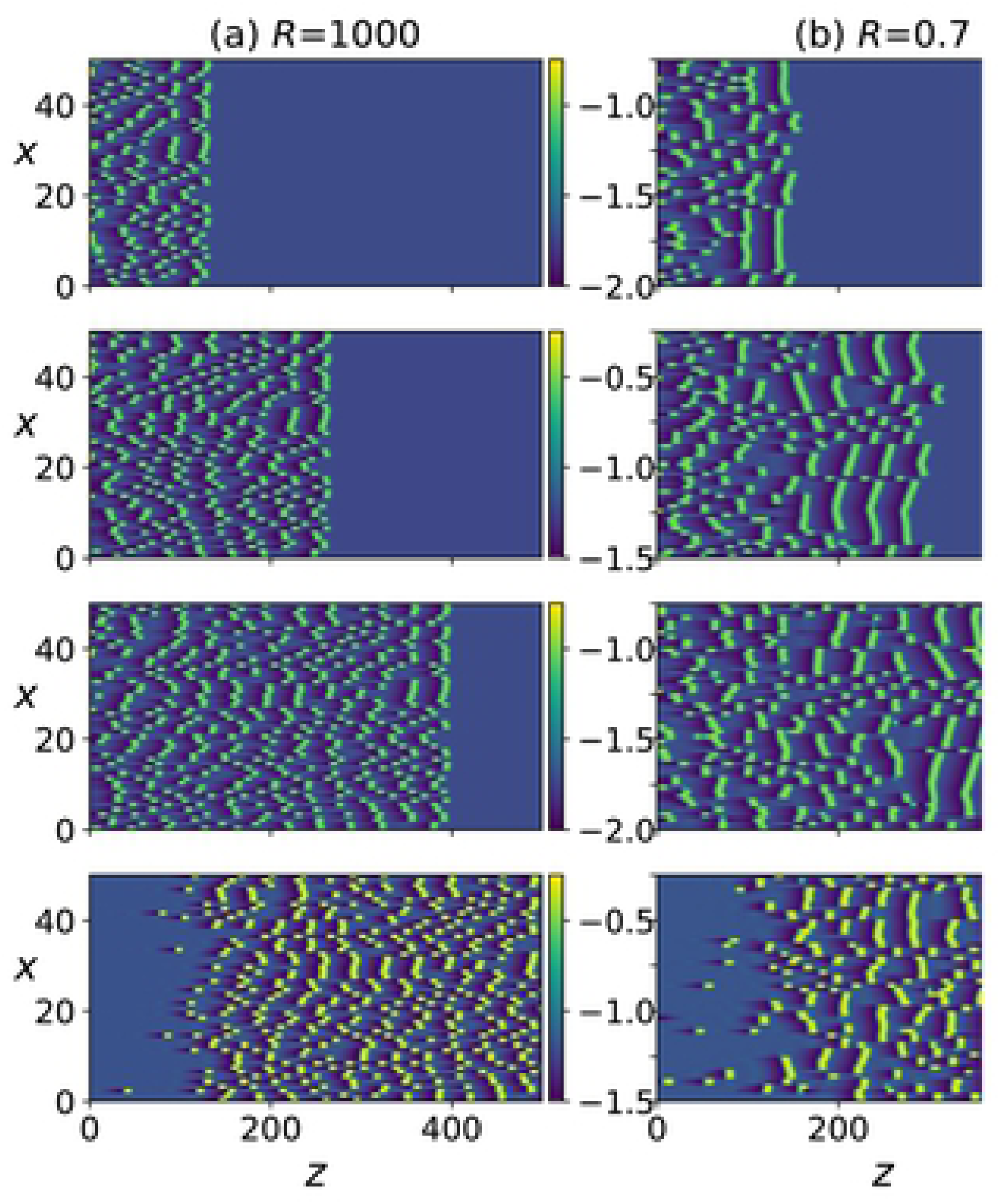
Numerical simulation of Eqs.13, the color bar indicates the value of *v*, the *x* variable indicates the axon number. (a) *R* = 0.8, axons number 30 and 20 are stimulated at *t* = 0 and *t* = 10, respectively and the panel rows from top to bottom correspond to *t* = 500, 1100, 1700, 2300, 2900. (b) same as in (a) but with axons number 25 and 24 stimulated. (c) same as in (b) but with stimulation at *t* = 0 and *t* = 11. (d) same as in (a) but with *R* = 0.4 and panels show *t* = 500, 1400, 2300, 3200, 4100.

### 2.2 Spatial patterns generation

Next, we increase the coupling strength by decreasing *R*, and observe that a transition occurs where each traveling impulse triggers new action potentials in its two immediately adjacent axons and the three neighboring impulses move together as a finite size traveling front, as shown in Fig.3(d). Further increase in coupling strength, leads to the next two adjacent axons being activated (Fig.4(a)).

**Fig 4.**
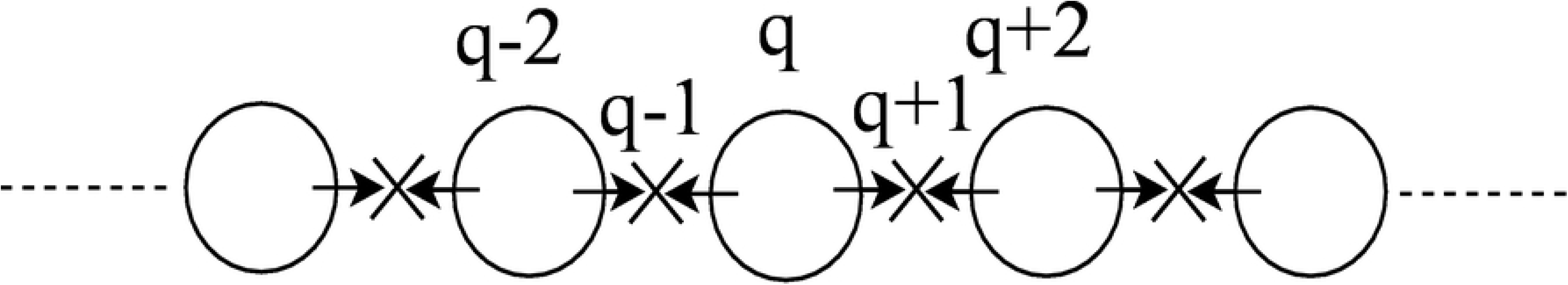
(a),(b) same as in Fig.3(a) but with *R* = 0.33 and *R* = 0.191, respectively. (c),(d) same as in Fig.3(d) but with *R* = 0.19 and *R* = 0.15, respectively.

Due to the presence of the scaling factor 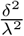 in the expression for *K*, we do not expect Eqs.15 to be equivalent to Eqs.13 for the same values of *R*. Nonetheless, it can be seen in the supporting information (S2 Fig) that the same behavior described so far also occurs in Eqs.15 as the coupling strength *K* is increased. In addition, as coupling strength is further increased, more and more impulses are triggered as the traveling front of impulses diffuses laterally and widens (Fig.5(a) and (b)). Even further increase in *K* (Figs.5(c) and (d)) leads to new fronts of impulses being induced in the forward but also backward direction, resulting in dynamic spatio-temporal patterns.

**Fig 5.**
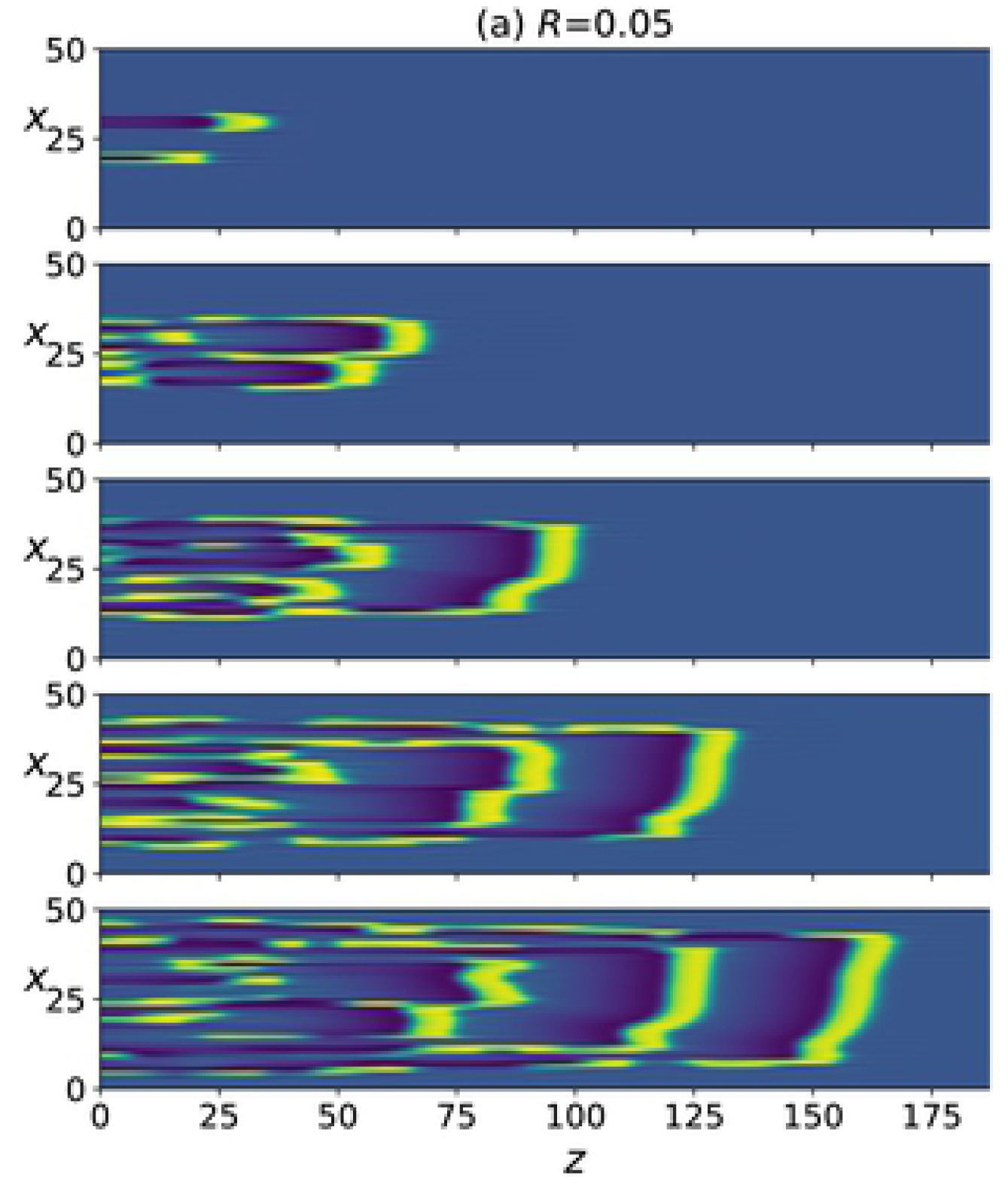
Same as in Fig.3 but for Eqs.15, with different values of *K* and for *t* = 500, 1300, 2200, 3100, 3999.

Figs.4(b),(c) and (d) show that the same transitions also occur in Eqs.13 as *R* is decreased, albeit the resulting patterns are more discrete and irregular. To better compare the responses of the two systems in this regime, we compare the discrete Fourier transform of the spatial patterns of the two systems at several time instants (supporting information S3 Fig). It can be seen that the spatial modal decomposition of the two is rather close, as quantified by the cosine similarity between the discrete Fourier transforms at every time step (supporting information S4 Fig). The mean value over time was ≈ 0.96 ± 0.006, which indicates close similarity. However, for Eqs.15, these spatio-temporal patterns persist only for a small range of the parameter, as a further increase in *K* causes the system to revert back to the laterally diffusing traveling front (Fig.6 right column). This is unlike Eqs.13 in which the complex spatio-temporal patterns persist as *R* is further decreased (Fig.6 left column).

**Fig 6.**
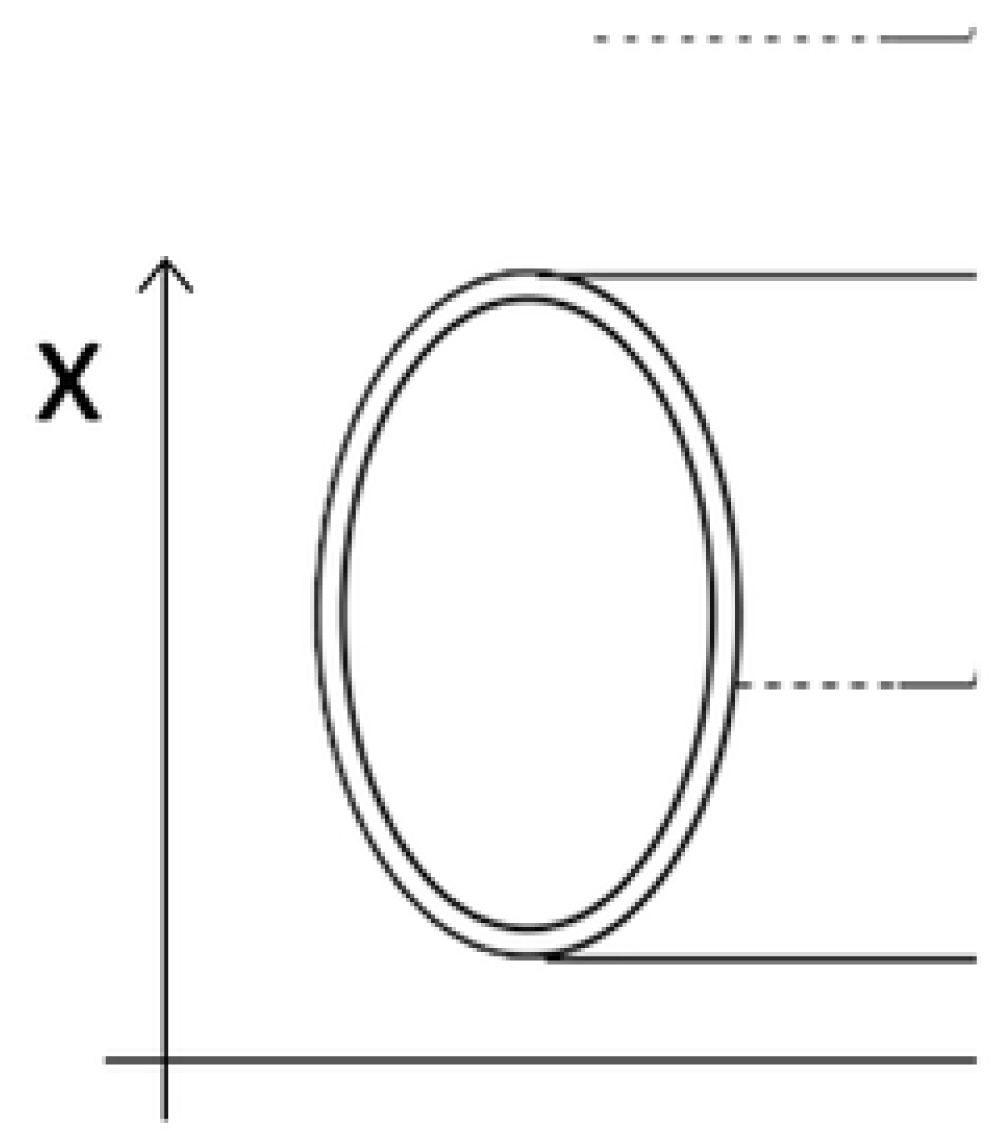
Numerical simulation of Eqs.13 for *R* = 0.05 (left) and of Eqs.15 for *K* = 0.05 (right). The panels from top to bottom correspond to *t* = 500, 1100, 1700, 2300, 2900.

### 2.3 Spiketrains interactions

Next, we stimulate all of the 50 axons and observe the collective dynamics. Each axon is stimulated by a finite train of 10 impulses. The intervals between impulses are generated by a Poisson process with a specified mean interval. The evolution of the resulting action potentials for the case of negligible ephaptic coupling is shown in Fig.7(a). When ephaptic coupling becomes significant, as in Fig.7(b), the impulses self-organize into phase locked traveling fronts. A similar effect also occurs for Eqs.15 (compare Fig.7 (c) and (d)).

**Fig 7.**
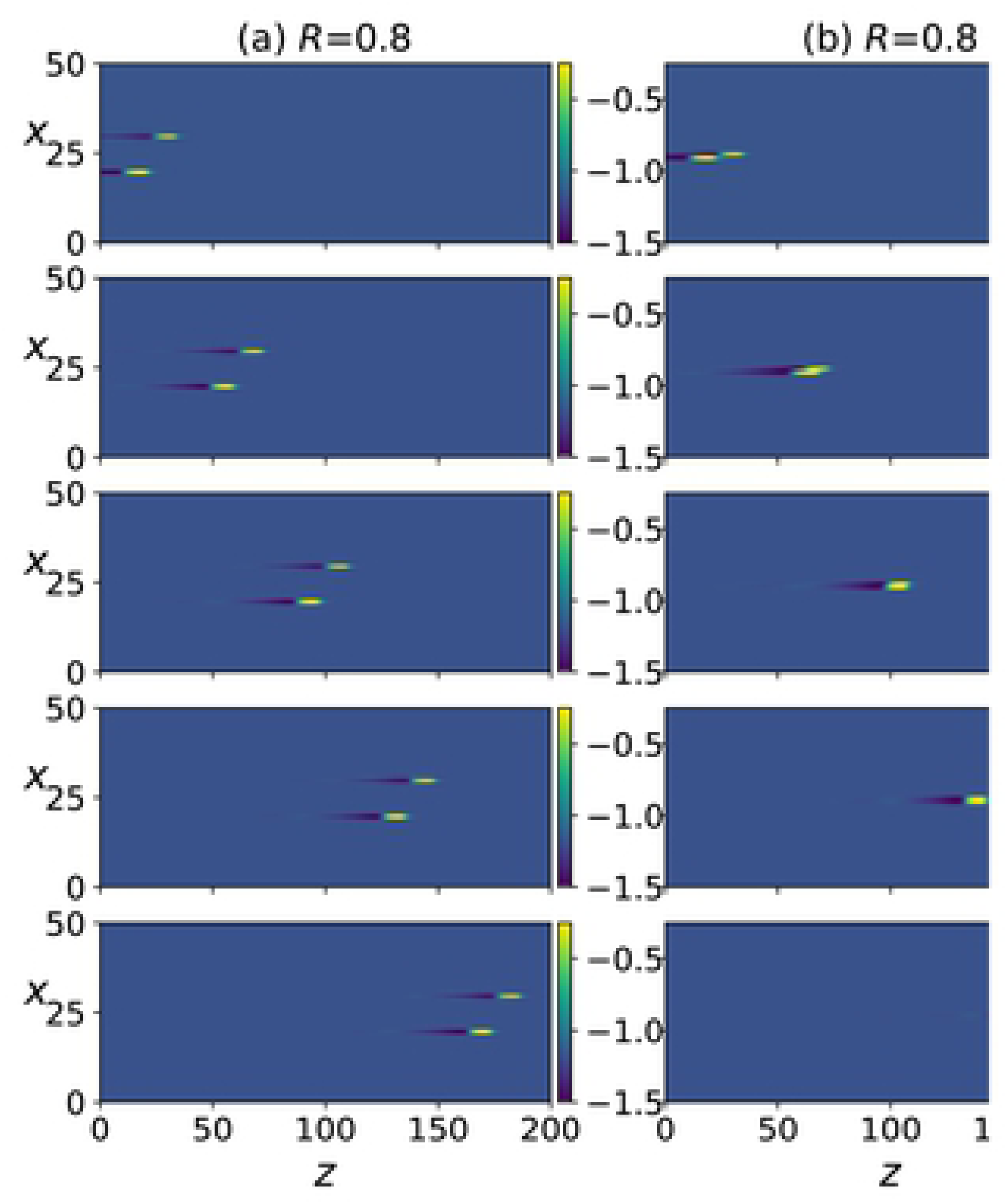
Numerical simulation of Eqs.13 ((a),(b)) and Eqs.15 ((c),(d)); spike trains of an average of 10 impulses are triggered along each axon, with a mean inter-impulse interval of 10. The panels from top to bottom correspond to *t* = 2499, 4999, 7499, 9999.

Previous work has shown that “in theory, whereas the pattern of Poisson-like impulse codes was modified during long-distance propagation, their mean rate was conserved” [39]. On the contrary, here in the presence of ephaptic interactions, the mean inter-spike interval (mISI), which is the inverse of the mean rate, decreases with increasing *z* location (supporting information S5 Fig and S6 Fig). The effect is clearly seen when the mISI is averaged over the 50 axons at downstream *z* locations (Fig.8). Note that Fig.7 shows only a portion of the simulation; in order to compute the mISI values shown in S5 Fig and S6 Fig, we let the simulation run long enough till all the impulses that were initiated at one end of the axon reach the other end, such that the total simulation time was 18000.

**Fig 8.**
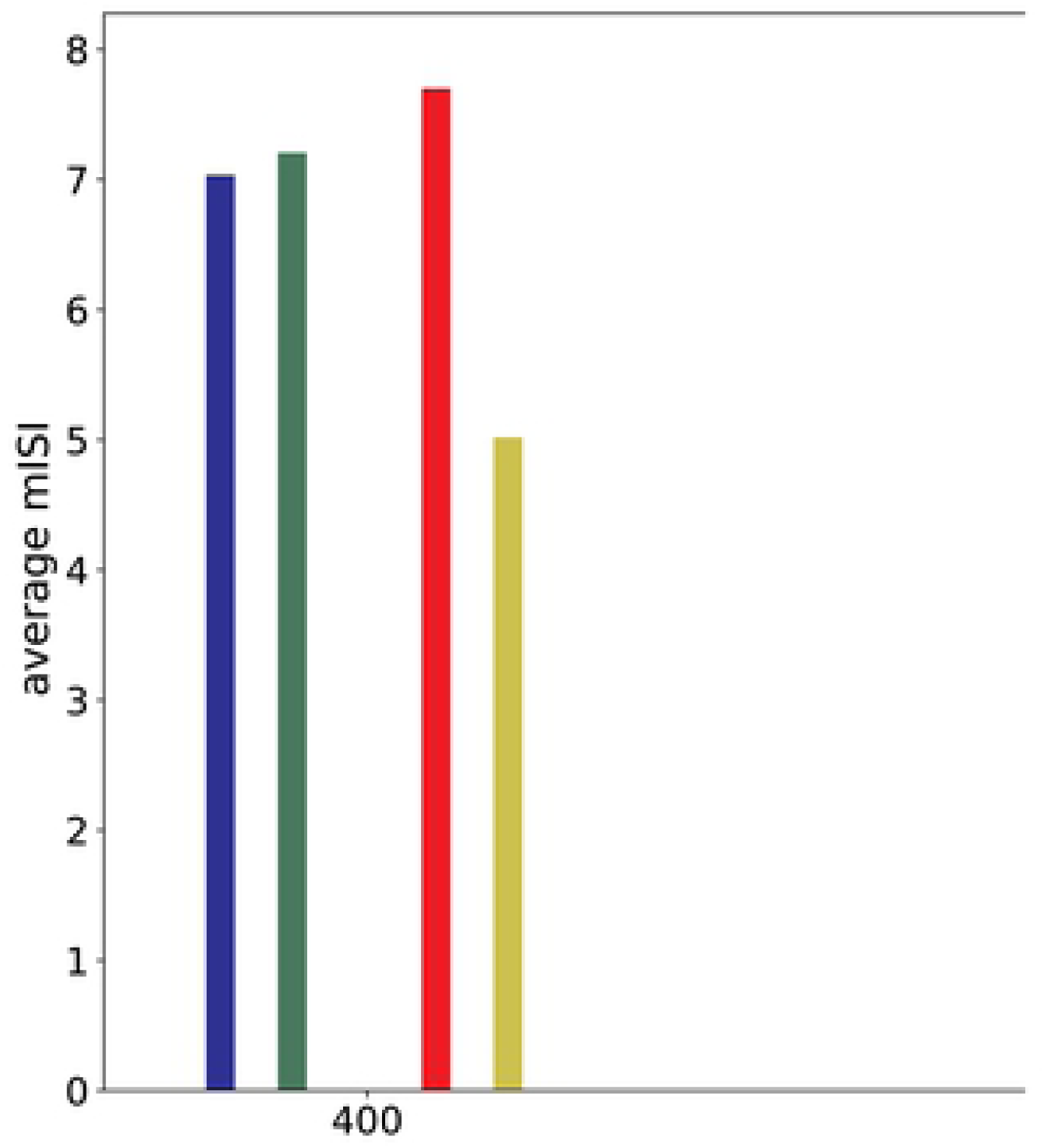
mISI averaged over the 50 axons for downstream *z* locations for the simulations in Fig.7(a) (blue), (b) (green), (c) (red) and (d) (yellow).

## 3 Discussion

We presented a minimal model for a sheet or volume of ephaptically coupled axons and explored its dynamics for a physically plausible range of parameters. We found that the model captures the experimentally observed attraction/repulsion effect between neighboring impulses. For strong enough coupling, the model predicts that action potentials traveling down an axon can trigger new action potentials in adjacent axons to be initiated and carried along with it forming a finite size traveling front. These fronts increase in size as more axons are recruited at higher coupling strength. Simulations with even higher coupling strength result in recurrence of impulses and backward propagation such that a pair of individual impulses initiated on two non-adjacent axons evolves into trains of impulses that diffuse laterally in the *x* direction as well as in both +ve and -ve *z* directions along the axons. We’ve also observed that ephaptic coupling can lead to self-organization among trains of impulses and significant alteration in the timing of action potentials which is known to be a key element in neuronal coding [40]. This suggests that ephaptic interactions along fiber pathways can theoretically play an active role in neuronal signal processing in the brain. The numerical simulations showed that the continuous limit approximation system mimics the qualitative behavior of the original model for a specific range of parameters. This continuous limit model offers the advantage of being mathematically more compact, more analytically tractable, less numerically expensive to solve and allows for easy extension of the model to full 3D geometry. It furthermore allows for a more intuitive interpretation of the ephaptic coupling terms and, in it’s integro-differential form, makes it intuitive that the ephaptic coupling creates a modulation of the diffusion in the axial direction with an alternating positive and negative diffusion term on a spatial length scale favoring structures on the scale of 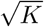.

In conclusion, we propose that the various nontrivial responses observed in our numerical exploration of ephaptic interaction might play an important and complex active role in inter-area neuronal signal transmission and processing in the brain. We hope these results will motivate a critical examination of the validity of the common assumption that neuronal fiber pathways merely act as transmission cables relaying signals between different brain regions. In contrast to that latter viewpoint, this theoretical investigation suggests that ephaptic interactions enhanced by the orientations and bundling of neuronal tracts in three-dimensional space can render the fiber pathways an active axonal medium that can give rise to complex spatio-temporal dynamics. If this emergent dynamics occurs under physiologically realistic conditions, then it would be a major so far unknown contributor to information processing in neural systems. We see various directions that this work can take in the future, including: further exploration of the rich repertoire of responses for different types of stimuli, accounting for variability in axonal diameters which will add spatial heterogeneity in the parameters, in addition, modifying the cable equation to include myelinated axons in the model. Finally, we hope that our work will inspire experimental work that can quantify and characterize the effects of ephaptic interactions in the axonal tracts of the brain.

## Supporting information

**S1 Fig. Shape of impulses along axons.** Two adjacent impulses starting out with small enough phase difference get attracted to each other and move in phase. Same as in Fig.3(b)

**S2 Fig. Numerical simulation results for Eqs.15 equivalent to results in Fig.3 for Eqs.13.** The color bar indicates the value of *v*, the *x* variable indicates the axon number. (a) *K* = 0.02, axons number 30 and 20 are stimulated at *t* = 0 and *t* = 10, respectively and the panel rows from top to bottom correspond to *t* = 500, 1100, 1700, 2300, 2900. (b) same as in (a) but with axons number 25 and 24 stimulated at *t* = 0 and *t* = 9, respectively. (c) same as in (b) but with stimulation at *t* = 0 and *t* = 10. (d) same as in (a) but with *K* = 0.025 and panels show *t* = 500, 1300, 2200, 3100, 3999.

**S3 Fig. Comparison of the spatial patterns of the two systems, Eqs.13 and Eqs.15.** Numerical simulation of Eqs.13 for *R* = 0.15 (left) and of Eqs.15 for *K* = 0.04 (right). The middle column shows a comparison of the spectrograms of the spatial discrete Fourier transform of the response of the two systems at specific times: Eqs.13 (red), Eqs.15 (blue). The panel rows from top to bottom correspond to *t* = 8000, 8600, 9350, 9950.

**S4 Fig. Cosine similarity between the discrete Fourier transform of the spatial patterns of the two systems.** The cosine similarity between the spatial discrete Fourier transform of the spatial patterns of the solutions of Eqs.13 and Eqs.15 with *R* = 0.15 and *K* = 0.04, respectively.

**S5 Fig. mISI for each axon at different *z* locations for Eqs.13.** For the simulation in Fig.7(a) *R* = 1000 (blue x) and (b) *R* = 0.7 (red dot).

**S6 Fig. mISI for each axon at different *z* locations for Eqs.15.** For the simulation in Fig.7(c) *K* = 0 (blue x) and (d) *K* = 0.025 (red dot).

